# Transient population dynamics drive the spread of invasive wild pigs in North America

**DOI:** 10.1101/2021.08.30.458238

**Authors:** Ryan S Miller, Michael A Tabak, Christopher L Burdett, David W Wolfson

**Author notes:** **Correspondence:** Ryan S. Miller, 2150 Centre Avenue, Bldg B, Fort Collins, Colorado, 80526 USA, office 1.970.215.2055 cell.

## Abstract

Invasion of nonindigenous species is considered one of the most urgent problems affecting native ecosystems and agricultural systems. Mechanistic models that account for short-term population dynamics can improve prediction because they incorporate differing demographic processes that link the environmental conditions of a spatial location explicitly with the invasion process. Yet short-term population dynamics are rarely accounted for in spatial models of invasive species spread.

Accounting for transient population dynamics, we predict the population growth rate and establishment probability of wild pigs following introduction into any location in North America. We compared predicted population growth rate with observed geographic rates of spread and found significant relationships between the annual rate of spread and population growth rates.

We used geospatial data on the distribution of mast producing tree species (a principle forage resource of wild pigs) and agricultural crops that can replace mast in their diets to predict population dynamics using transient population simulations. We simulated populations under different initial population sizes (i.e. number of introduced individuals, often termed propagule size) and for different amounts of time following introduction. By varying the initial population size and simulation time, we were able to identify areas in North America with high probability for establishment of wild pigs if introduced. Our findings can be used to inform surveillance and removal efforts to reduce the potential for establishment and spread of wild pigs.

## Introduction

Invasion of nonindigenous species is considered one of the most urgent problems affecting native ecosystems with 87% of imperiled species threatened by invasive species (Mack et al., 2000; McClure et al., 2018). Additionally, nonindigenous species threaten agricultural systems requiring considerable policy activity to mitigate invasive species concerns (Miller et al., 2018; Miller, 2020). Existing theory indicates that the distribution and spread of invasive species is the result of complex ecological processes that include the frequency and size of introduction (propagule pressure), species-specific traits that provide a fitness advantage (high reproductive capacity and dispersal), and biotic and abiotic characteristics of the recipient ecosystem that limit or facilitate invasion (Simberloff, 2009; Lustig et al., 2017). However dynamic risk assessments of establishment and spread typically lack explicit consideration of these interacting factors (Catford et al., 2011; Gallien et al., 2015) and broad scale projections of invasive species distributions are typically based on static approaches linking species occurrence to biotic and abiotic factors (Guisan and Zimmermann, 2000; Lustig et al., 2017). Mechanistic models that account for short-term population dynamics can improve prediction because they incorporate differing demographic processes that link the environmental conditions of a spatial location explicitly with the invasion process. Yet short-term population dynamics are rarely accounted for in spatial models of invasive species spread (Lustig et al., 2017).

When populations are not at equilibrium (e.g., the age structure is not the stable age structure and population growth are not defined by the equilibrium population growth rate, λ), they are expected to exhibit transient dynamics (Caswell and Werner, 1978; Hodgson and Townley, 2004; Tremblay et al., 2015). Transient dynamics can cause populations either grow or shrink at a much faster rate than would be expected under equilibrium conditions (Stott et al., 2011; McDonald et al., 2017). When a small group of conspecific individuals is introduced to a new location (a “propagule”), it is not likely to be a stable population, and transient dynamics can lead to either rapid growth and establishment of a population or local extinction (Iles et al., 2016).

Invasive wild pigs (*Sus scrofa*), also known as feral hogs, feral swine, or wild boar, are recognized as one of the most widespread and destructive invasive species in the world (Barrios-Garcia and Ballari, 2012). Wild pigs are native to Eurasia and Northern Africa, but have been widely introduced for centuries, often deliberately, and now occupy every continent except Antarctica (Mayer and Brisbin, 1991; Lewis et al., 2017). In North America it is an extremely destructive invasive species that is of concern for human and animal health (Miller et al., 2017), causes significant economic damage (Anderson et al., 2016), negatively impacts imperiled species (McClure et al., 2018), and is commonly introduced into new locations (Tabak et al., 2016; Hernandez et al., 2018). These concerns have generated significant Federal policy to mitigate these impacts with the US Department of Agriculture establishing the Animal Plant Health Inspection Service (APHIS) National Feral Swine Damage Management Program in 2013 (federal government fiscal year 2014) aimed at reducing the spread of wild pigs in the United States (USDA, 2015; Miller et al., 2018). Nevertheless, the distribution of wild pigs in Canada and the United States has increased dramatically in recent years (Michel et al., 2017; Snow et al., 2017), and research suggests that a major mechanism for their spread is translocation of individuals to augment populations for recreational hunting (Tabak et al., 2016). Wild pigs evolved as pulsed resource consumers of mast crops; they have larger litter sizes and reproduce more often under favorable forage conditions, and have reduced fecundity under poor forage conditions (Ostfeld and Keesing, 2000). This elasticity in fecundity makes wild pigs especially susceptible to transient dynamics (Bieber and Ruf, 2005; Tabak et al., 2018). Despite their ancestral dependence on mast crops, wild pigs have evolved to be dietary generalists, as agricultural crops can replace mast in their diets (Schley and Roper, 2003; Rosell et al., 2012) and they currently thrive in ecosystems that lack mast producing species (Caley, 1997; Choquenot and Ruscoe, 2003). Nevertheless, their reproductive biology retains this elasticity of fecundity, which can cause populations to rapidly grow and establish following introduction in new environments, leading to the further expansion of this species. Despite significant efforts to forecast the spread of wild pigs (Snow et al., 2017), estimate the probability of occurrence (McClure et al., 2015), and predict the potential density of wild pig populations (Lewis et al., 2017) there are no available spatial predictions of population growth and establishment risk if wild pigs were released into a given spatial location.

Our objective was to predict the population growth rate and establishment probability of wild pigs (accounting for transient dynamics) following introduction into any location in North America. We then compared predicted population growth rate with observed geographic rates of spread to determine if increased population growth was associated with increased spread (e.g. invasion). We used geospatial data on the distribution of mast producing tree species (a principle forage resource of wild pigs) and agricultural crops that can replace mast in their diets to predict population dynamics using transient population simulations. We simulated populations under different initial population sizes (i.e. number of introduced individuals, often termed propagule size) and for different amounts of time following introduction. By varying the initial population size and simulation time, we were able to identify areas in North America with high probability for establishment of wild pigs if introduced. Our findings can be used to inform proactive surveillance and removal efforts to reduce the potential for establishment and spread of wild pigs.

## Methods

### Predicting population growth and establishment

We used transient population dynamics models to simulate population trajectories for invasive wild pig populations under different types of environments that they could potentially experience in North America. Survival and fecundity rates for different mast qualities (poor, intermediate, and good) were obtained from the literature (Briedermann, 1967; Bieber and Ruf, 2005). Data are scarcely available for the quality of mast from trees over time. In the literature we found historic data from one common mast species, *Fagus sylvatica*, over 114 years (Hilton and Packham, 2003) and from a community of five mast tree species, *Quercus* spp., over 12 years (Koenig et al., 1994). Following Tabak et al. (2018), we used these historic mast data to simulate three broad types of environmental conditions to which wild pigs might be introduced: an environment with one mast tree species, an environment with a mast community (containing five mast tree species), and an environment with agricultural subsidy. We conducted simulations in these three environments under nine different introduction scenarios: we used propagule sizes of 5, 10, or 20 as the number of females introduced at the beginning of the simulations, and for each of these propagule sizes, we allowed simulations to run for 1, 5, or 10 years.

We estimated stochastic population growth rate (*λ*_*s*_) and probability of establishment for wild pig populations in these environments and in environments with agricultural subsidy using the methods and the R scripts of Tabak et al. (2018). A simulated population was determined to have established if *λ*_*s*_ > 1 and population size at the end of simulation was ≥ 60. Tabak et al. (2018) provide justification for using 60 individuals as the threshold for population establishment in wild pig populations. Unfortunately, data are unavailable for mast records from areas with 2-4 mast species and from areas with > 5 mast species. Therefore, we assumed that five mast species represented the maximum population growth potential resulting from their consumption of mast crops. For areas with 2-4 mast species, we estimated *λ*_*s*_ and establishment probability using loess regression, where the predictor variable was the number of mast species (1 or 5) and the response variable was *λ*_*s*_ or establishment probability. To calculate the *λ*_*s*_ in areas with both mast and agriculture, we estimated *λ*_*s*_ as the mean between the growth rate estimated as a result of mast and the growth rate estimated as a result of agricultural subsidy. To calculate establishment probability in such situations, we took the sum of probability of establishment that was estimated as a result of mast species and that resulting from agricultural subsidy.

There are clearly limitations in our estimation of growth rate and of establishment probability as we extrapolated to areas with numbers of mast trees for which we do not know the frequency of mast and we are unaware of how wild pig growth potential changes in the presence of both agriculture and mast trees. We also only considered the effects of forage on population growth, but other factors might affect growth rate (e.g., climate could have strong effects on survival). However, we are interested in the general trends of population growth and establishment across the continent more than precisely estimating these rates.

To apply these growth rates and establishment probabilities to geographic locations, we used a geospatial layer depicting the number of mast species in each 1 km^2^ (Burdett et al. unpublished data) and the presence of crops that can replace mast in wild pigs’ diets using the USDA’s cropscape data (NASS, 2019). Cropscape data are only available for USA, and we wanted to extend our model results to all of North America, so we also used the consensus land□cover layer for North America (Tuanmu and Jetz, 2014) and compared this with the results in USA that were produced using the detailed cropscape data (Fig S3). Since the maps created using these two datasets were similar for the USA, we assume that it is reasonable to use the consensus land-cover and apply our results to all of North America.

### Determining rates of spread

We estimated the geographic rate of spread of wild pigs in the contiguous U.S. using data from the National Feral Swine Mapping System (Corn and Jordan, 2017). These data describing wild pig distribution were compiled at irregular intervals from 1982-2008 and annually since 2008. They are the best available data describing the known distribution of wild pigs over the past 36 years and have been used to forecast the spread of wild pigs (Snow et al. 2017), estimate the probability of occurrence (McClure et al. 2015), determine wild pig agricultural damage risk (Miller et al. 2017), predict federal policy to control wild pigs (Miller et al. 2018), and determine the risk wild pigs pose to imperiled species (McClure et al. 2018). Polygons representing the known geographic extent of established wild pig populations (defined as populations present for two or more years with evidence of reproduction) are reported to the National Feral Swine Mapping System nationally by wildlife professionals in state wildlife resource agencies and the United States Department of Agriculture. The resulting distribution data is curated for accuracy with national data available annually.

Polygons representing the observed distribution for 1982, 1988, 2004, 2008, 2013, and 2017 were aggregated to watersheds (HUC12) as described in McClure et al. (2015) to discretize consistent, comparable, and ecologically relevant sampling units. The annual watershed rate of spread among HUC12 watersheds was then calculated for three courser watershed scales – HUC4, HUC6, HUC8. This was done because the spatial scale used for analysis can influence inference (Farnsworth et al. 2006). We calculated the annual rate of spread (*θ*) for each watershed *j* in each time period *t* as,

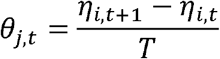

Where *η_i,t_* is the count of HUC12 watersheds occupied by wild pigs in the coarser watershed *j* in year *t*, and *T* is the number of years between t and t+ 1. We also calculated the mean annual rate of spread from 1982 to 2013, the final year in this range was the beginning of a National Feral Swine Damage Management Program to control wild pigs, and from 2013 to 2017 which represents the period after the National program began.

To determine if the predicted population growth rate was associated with the annual rate of spread (*θ*) in watersheds we regressed the annual rate of spread, *θ*, on the mean watershed population growth rate, *λ*_*s*_. This was implemented using a linear model assuming a Gaussian error structure. We used adjusted R^2^ as a measure of goodness of fit and the predictive capacity of each model (Kutner et al., 2005).

## Results

Our spatial estimates of *λ*_*s*_ that used the detailed cropscape data for the US were similar to estimates using the cultivated layer that is available for all of North America (Fig. 1). Since these estimates were similar and using the cultivated data allows our results to have broader interpretation, we focus on the continental scale for the remainder of this article. With a small initial population size (propagule size = 5), mean *λ*_*s*_ was < 1, regardless of the number of years in which we allowed simulations to run following introduction (Fig. 1). As we increased propagule size and the number of years of simulation, *λ*_*s*_ increased so that with a propagule size of 20 and ten years of simulation, *λ*_*s*_ was > 1 for almost all of North America. We found the highest growth rates in the upper Midwestern United States and the southern prairie region of Canada.

**Figure 1.**
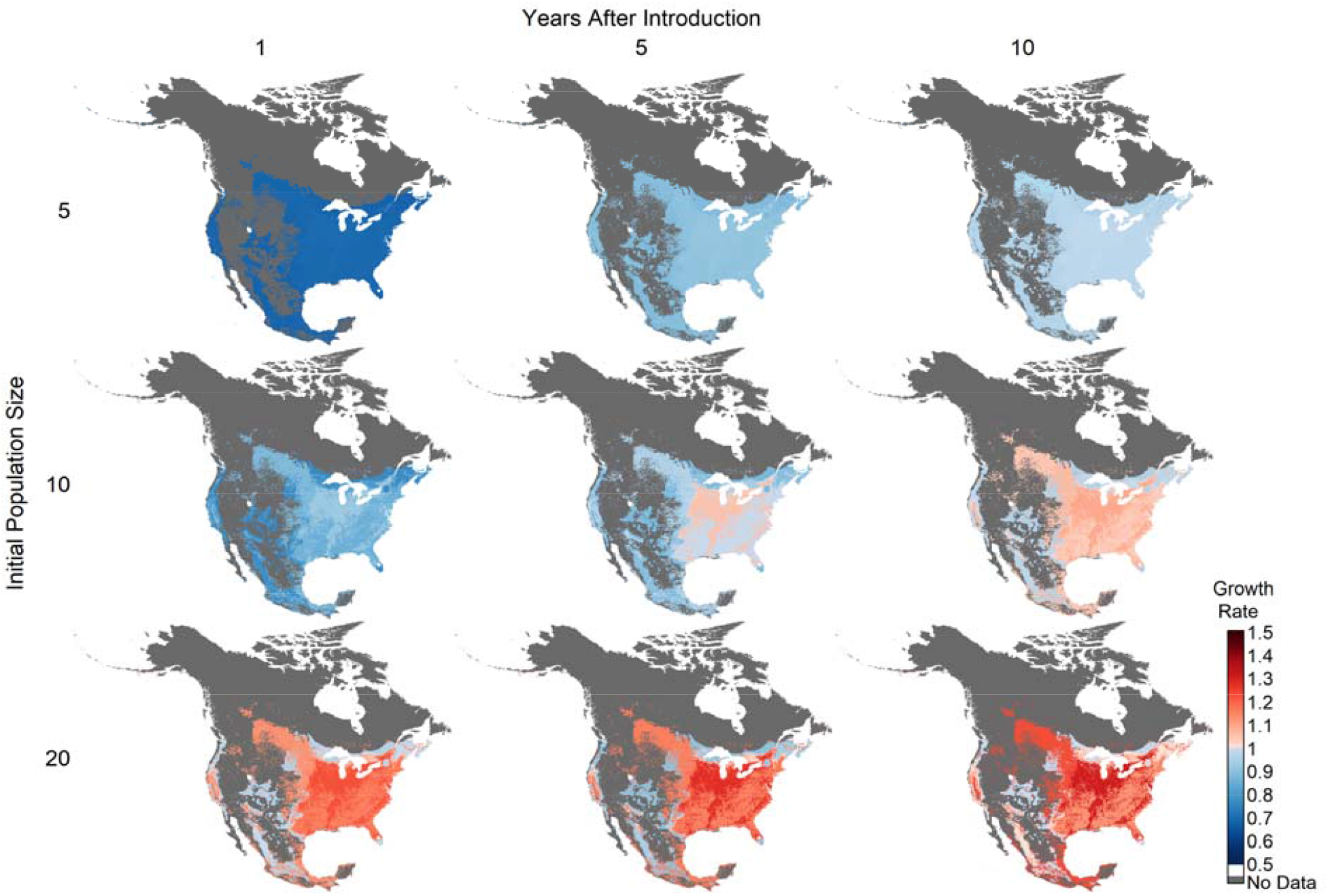
Estimated transient population growth rates (*λ*_*s*_) for wild pigs in North America by years after introduction (x axis) and initial population size (y axis).

The mean establishment probability was very low (< 6%) when the initial population size was 5 females (Fig. 2). When propagule size was increased to 20 and simulations ran for 5 to 10 years, establishment probabilities were high in the same locations we found the highest growth rates (Figure 1 and 2). Establishment probabilities approached 1 for many locations in the upper Midwestern US.

**Figure 2.**
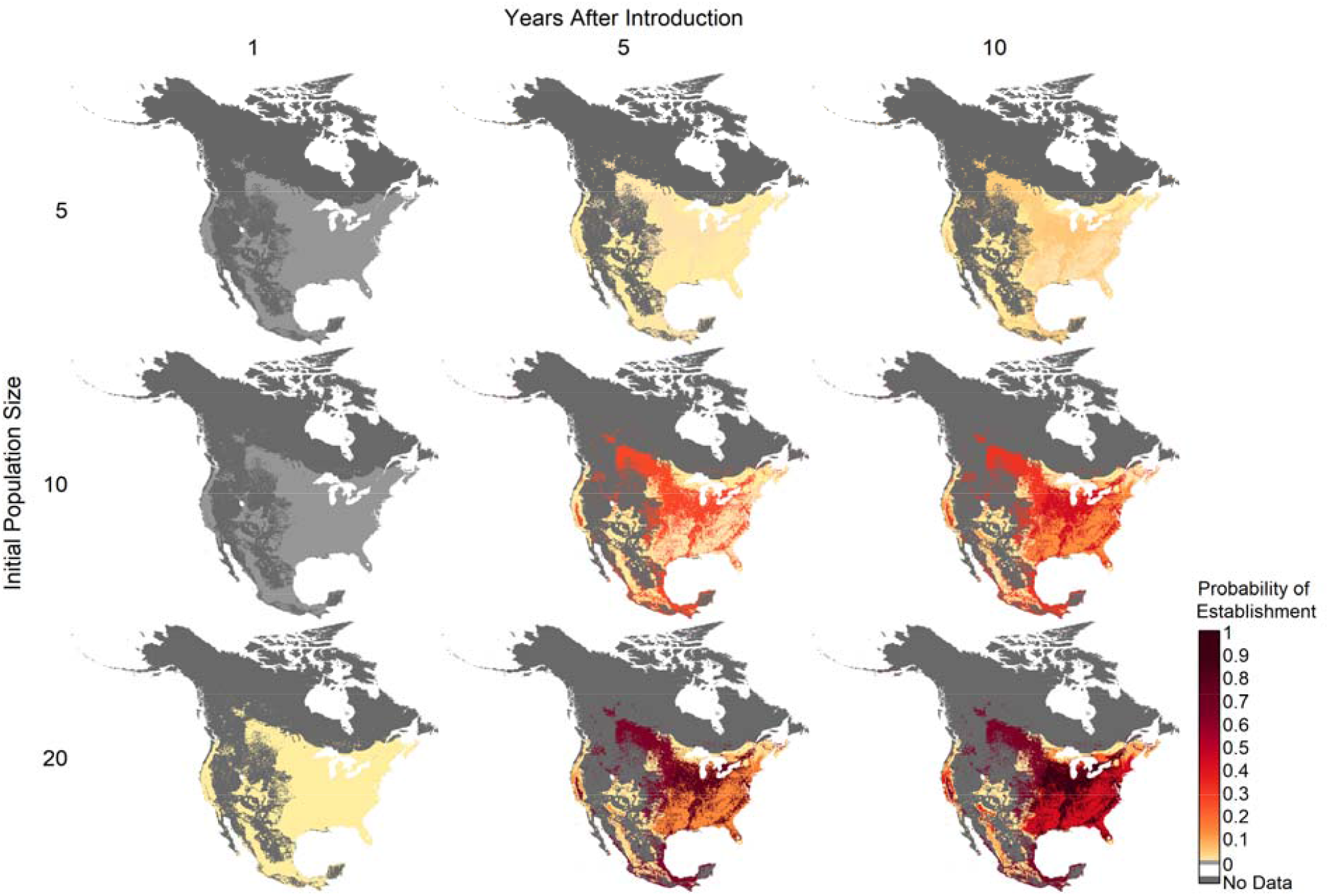
Estimated probability of establishment for wild pigs in North America by years after introduction (x axis) and initial population size (y axis).

### Annual spread rate

The annual spread rate (θ) was proportionally consistent across the three watershed scales however it demonstrated strong spatial heterogeneity (Figure 3, see supplemental for all watershed scales). Mean annual rates of spread changed among time periods with the greatest rates of spread occurring from 1988 to 2004 across all watershed scales (θ_*HUC4*_ =4.54, θ_*HUC6*_ =2.86, θ_*HUCS*_ =0.65) (see supplemental for all spread rates). Annual rates of spread demonstrated spatial heterogeneity and were not constantly greatest along the northern extent of wild pig distribution. During the earliest periods from 1982 to 2004 spread rates were positive across the majority of the wild pig distribution. From 2004 to 2013 watersheds along the Red River, Rio Grande River, and Ohio River consistently had the highest spread rates. Annual spread rates declined significantly across all watershed scales after the establishment of the National Feral Swine control program in 2013 with some regions having negative spread rates. The highest rates of spread after 2013 where confined to the Ohio River and Tennessee River watersheds.

**Figure 3.**
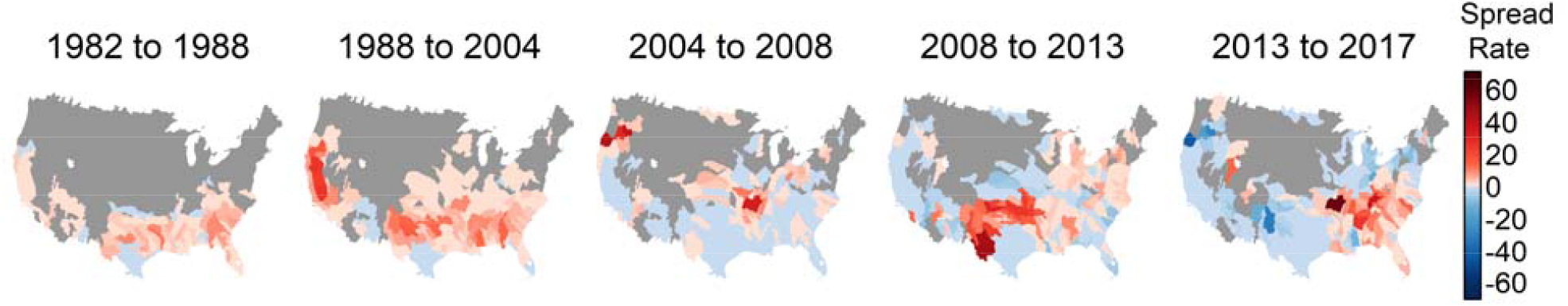
Annual watershed level spread rate (watersheds/year) for wild pigs from 1982 to 2017 for three watershed scales. Red indicates positive rates of spread and blue indicates negative rates of spread (i.e. contractions in the number of occupied watersheds).

### Association of spread rate and population growth rate

We found significant relationships between the annual rate of spread and population growth rates. From 1982 to 2013, the period prior to the National Feral Swine control program, population growth rates were a significant positive predictor of the annual rate of spread for all watershed scales (Fig. 4). The mean population growth rate explained a large amount of the variation in rate of spread with adjusted R^2^ values ranging from 0.729 to 0.953 for the periods from 1982 to 2013 (see supplemental Table S2). Additionally, the effect of population growth rates on spread rates increased 47.6% (HUC4), 53.2% (HUC6) and 54.6% (HUC8) from the earliest period (1982 to 1988) to the period just prior to the program (2008 to 2013). For the period after the establishment of the Feral Swine control program (2013 to 2017), population growth rates were a significant positive predictor of the annual rate of spread only at the HUC8 watershed scale.

**Figure 4.**
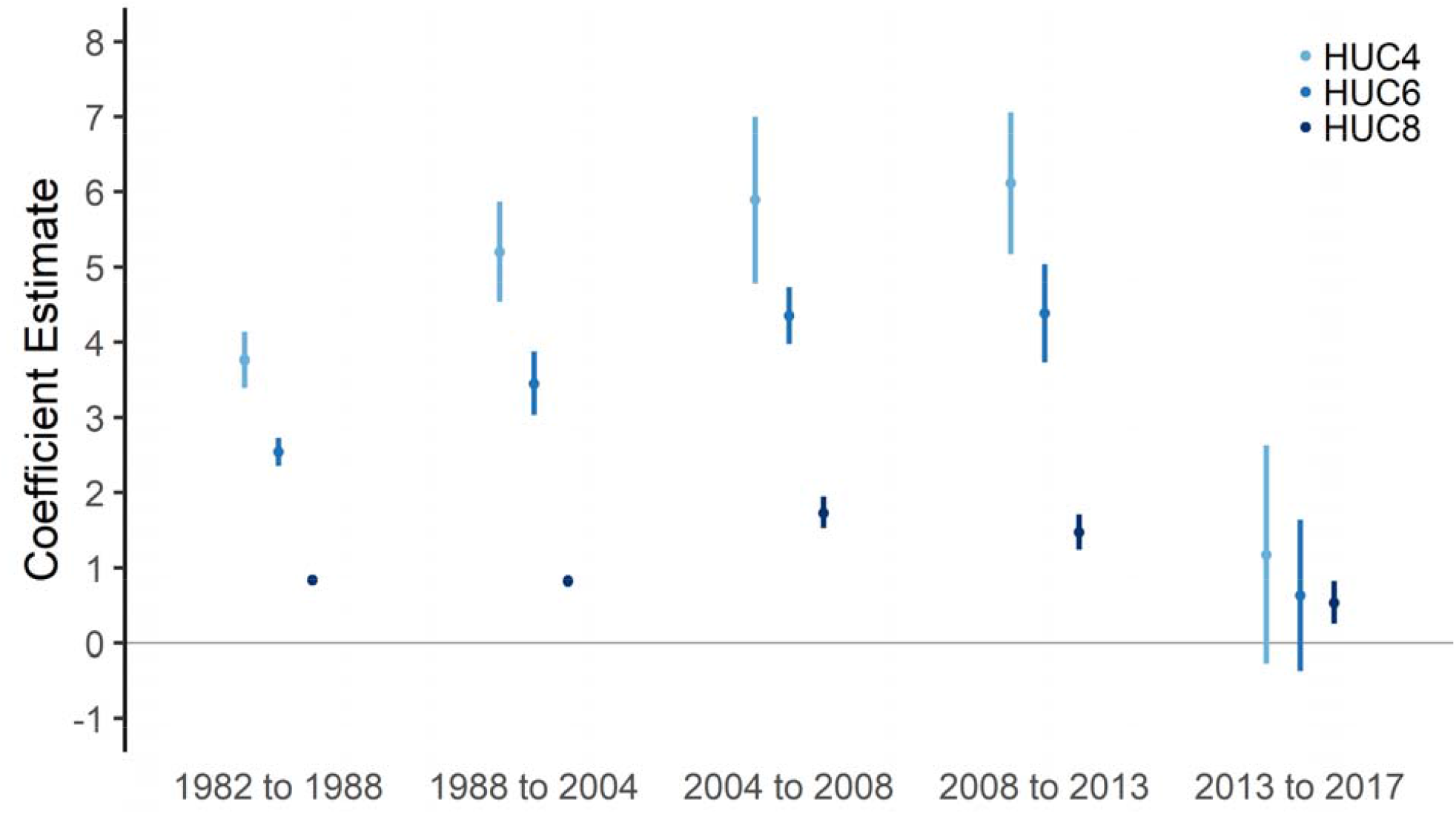
Regression coefficients for linear models regressing wild pig spread rate on transient population growth rate. Spread rate was increasingly positively associated with transient population growth rate in all time periods except the period from 2013 to 2017. A national wild pig control program was initiated beginning in 2013.

## Discussion

Explicitly accounting for initial population size and age structure allowed us to account for transient dynamics and demographic stochasticity that influence predictions of population growth and probability of establishment. Our geospatial projections of population growth rate and establishment probability (Figs. 2 and 3) highlight areas at increased risk of successful establishment of wild pig populations if introduced. The strong correlation between our estimates of population growth (*λ*_*s*_) and the annual watershed rate of spread (*θ*) supports the validity of our population growth estimates. In other systems, transient population growth rate has been identified as important in predicting establishment success and long term viability of invading species (Iles et al., 2016). Our results can be used to target areas for surveillance that have higher potential for wild pig establishment and rapid population growth allowing for early identification of wild pigs, as well as efforts to minimize population expansion once new wild pig populations are identified.

### Applying mechanistic models to geospatial projections

While mechanistic models are often used for distribution and niche modeling for invasive species (Kearney and Porter, 2009; Peterson et al., 2015), population dynamic processes across spatial scales are more commonly evaluated using statistical models to correlate population processes with spatial covariates (but see Chandler et al., 2018; Quintana□Ascencio et al., 2018). Correlative spatial models are convenient, especially in systems with less biological data, because they do not require an understanding of the mechanistic links between an organism and its environment. However, mechanistic processes tend to limit the expansion of species distributions, so it can be useful to build models that evaluate mechanistic processes across large spatial extents when the goal is to predict the growth and establishment potential of a species. Furthermore, for populations that experience transient dynamics, mechanistic models may be more applicable than correlative statistical models. When a population is introduced into a new environment, it will usually exhibit transient dynamics (Iles et al., 2016) and wild pigs are especially prone to transient dynamics due to their evolution as pulsed resource consumers (Bieber and Ruf, 2005; Tabak et al., 2018).

### Comparison to other studies

Our estimates of population growth and establishment probability correspond well to the reported establishment of wild pigs throughout the United States and the prairie provinces of central Canada (Michel et al., 2017). However, our estimates of establishment probability differ in several ways from the predicted spread of wild pigs in the United States (Snow et al., 2017). We found high establishment probability throughout the upper Mississippi and Ohio River drainages while previous studies predicted lower probability of spread in these areas. The historical spread of wild pigs in the United States by Snow et al. (2017) implicitly includes factors such as human translocation of wild pigs which previous studies have found to be important in the spread of wild pigs (Tabak et al., 2016). The differences between Snow et al. (2017) and our results may stem from differences in the frequency of introduction, which is likely driven by anthropocentric factors. This indicates that while the probability of establishment is generally high in some regions of the U.S., the historical frequency of introduction has likely been low. Our predicted probability of establishment and population growth rates generally correspond well with the predicted equilibrium population density of wild pigs by (Lewis et al., 2017).

The differences among the predictions of wild pig spread (Snow et al., 2017), population density (Lewis et al., 2017), and probability of establishment (McClure et al., 2015) are not mutually exclusive. The differences among these results is largely due to each study focusing on a different component of the invasion process-introduction, establishment, population growth, and spread (Reise et al., 2006). These differences highlight the need to develop integrated mechanistic approaches that explicitly include factors that drive introduction pressure (e.g. Snow et al. (2017) and Tabak et al. (2016)) probability of establishment and population growth (e.g. our study), and factors regulating long-term population growth and equilibrium population size (e.g. Lewis et al., 2017). Accounting for the invasion process in a spatially explicit context would allow for greatly improved predictions of establishment risk and consequences once established.

### Management implications

Establishment probability was very low when initial population size was small and when populations had fewer years to become established (Fig. 3), because populations in these simulations were unlikely to meet one of the criteria for establishment: a population size of 60 individuals in the final simulation year. This indicates that there may be the opportunity to prevent population establishment if actions are undertaken quickly upon recognizing newly introduced populations. With larger initial populations and more time following introduction, the potential for population establishment increased dramatically, indicating that it is best to prevent introductions of large numbers of individuals and to begin interventions early. Reducing the frequency and size (number of animals) of releases of wild pigs can reduce establishment probability, but these goals may be difficult to implement, as movement and release of wild pigs is common and has been documented on multiple continents (Hampton et al., 2004; Tabak et al., 2016; Hernandez et al., 2018). This is often complicated by limitations of current tools available for detecting newly released pigs. Tools such as pig detection dogs (Keiter et al., 2016) and passive monitoring using camera traps (Tabak et al., 2019) are difficult to deploy over large geographic extents for long periods of time. Our results can be used to refine where monitoring using these tools is conducted, improving the chances of early detection of pigs. When considering management to prevent population establishment, it is important to note that our methods only account for a single introduction event so likely represent a conservative estimate of establishment probability. Wild pigs are often repeatedly introduced to the same location (Mayer and Brisbin, 1991), which can increase establishment probability through demographic rescue of the founder population (Hufbauer et al., 2015).

### Extensions

Our method of estimating population growth and establishment success has some important limitations that offer opportunities to extend our approach. Complete data on the frequency and intensity of mast and how this changes with the size and community structure of mast species is limited or not available. To address this data limitation we extrapolated using available data, however there is uncertainty in our approach for making this extrapolation. The development of data describing mast frequency and intensity over large geographies in North America would have significant value for population dynamic modeling of many species that consume seasonal mast and fruit. Additionally, we used data linking how mast frequency and intensity affect vital rates of wild pigs from Europe (Briedermann, 1967; Bieber and Ruf, 2005); data describing this relationship in other continents are needed to better extrapolate globally. In our predictions the primary driver of wild pig vital rates was assumed to be access to mast. However, previous studies have indicated that precipitation and temperature can influence survival of younger age classes impacting transient population dynamics of wild pigs (Miller, 2017). Currently most studies reporting survival, litter size, and frequency of litters and how these factors correlate with environmental conditions are from wild pig populations in Europe. Studies collecting these vital rates for wild pig populations in North American would be of great value. We also do not account for inter and intra species interactions that may also be limiting for founder populations. For example Lewis et al. (2017) found that predator species richness was negatively associated with wild pig population density. Accounting for these interactions within a dynamical modeling framework that accounts for transient dynamics would be an exciting extension of our work.

Despite these limitations, predicted population growth and the spread rate of wild pigs in areas where they have been introduced were highly correlated. The positive correlation indicates that despite the data uncertainties, our projections are reasonable estimates of population growth and establishment potential for wild pigs.

### Conclusion

Risk assessments of establishment and spread of invasive species that consider the frequency and size of introduction along with the biotic and abiotic characteristics of the recipient ecosystem using mechanistic models that account for short-term population dynamics can improve prediction. Accounting for short-term population dynamics in spatial models of invasive species spread will become increasingly important as variation in climatic conditions increases. For many invasive species, including wild pigs, increasing temperature is expected to increase invasion success and expand regions with the potential for invasion (Vetter et al., 2015). Spatial models that account for short-term population dynamics can allow for uncertainties resulting from increased climate variation to be explicitly considered, improving predictions. An important extension of our work is to integrate, using mechanistic approaches that account for short-term population dynamics, with ecological and anthropocentric processes that determine introduction, establishment, and spread of invasive species.

## References

Anderson, A., C. Slootmaker, E. Harper, J. Holderieath, and S. A. Shwiff. 2016. Economic estimates of feral swine damage and control in 11 US states. Crop Protection 89:89–94.

Barrios-Garcia, M. N., and S. A. Ballari. 2012. Impact of wild boar (Sus scrofa) in its introduced and native range: a review. Biological Invasions 14(11):2283–2300.

Bieber, C., and T. Ruf. 2005. Population dynamics in wild boar Sus scrofa: ecology, elasticity of growth rate and implications for the management of pulsed resource consumers. Journal of Applied Ecology 42(6):1203–1213.

Briedermann, L. 1967. Studies on the diet of wild boar in lowland of the German Democratic Republic, German Academy of Agricultural Sciences, Berlin.

Caley, P. 1997. Movements, activity patterns and habitat use of feral pigs (Sus scrofa) in a tropical habitat. Wildlife Research 24(1):77–87.

Caswell, H., and P. A. Werner. 1978. Transient behavior and life history analysis of teasel (Dipsacus sylvestris Huds.). Ecology 59(1):53–66.

Catford, J. A., B. J. Downes, C. J. Gippel, and P. A. Vesk. 2011. Flow regulation reduces native plant cover and facilitates exotic invasion in riparian wetlands. Journal of Applied Ecology 48(2):432–442.

Chandler, R. B., J. Hepinstall-Cymerman, S. Merker, H. Abernathy-Conners, and R. J. Cooper. 2018. Characterizing spatio-temporal variation in survival and recruitment with integrated population models. The Auk: Ornithological Advances 135(3):409–426.

Choquenot, D., and W. A. Ruscoe. 2003. Landscape complementation and food limitation of large herbivores: habitat-related constraints on the foraging efficiency of wild pigs. Journal of Animal Ecology 72(1):14–26.

Corn, J. L., and T. R. Jordan. 2017. Development of the national feral swine map, 1982–2016. Wildlife Society Bulletin 41(4):758–763.

Gallien, L., F. Mazel, S. Lavergne, J. Renaud, R. Douzet, and W. Thuiller. 2015. Contrasting the effects of environment, dispersal and biotic interactions to explain the distribution of invasive plants in alpine communities. Biological invasions 17(5):1407–1423.

Guisan, A., and N. E. Zimmermann. 2000. Predictive habitat distribution models in ecology. Ecological modelling 135(2-3):147–186.

Hampton, J. O., P. B. Spencer, D. L. Alpers, L. E. Twigg, A. P. Woolnough, J. Doust, T. Higgs, and J. Pluske. 2004. Molecular techniques, wildlife management and the importance of genetic population structure and dispersal: a case study with feral pigs. Journal of Applied Ecology 41(4):735–743.

Hernandez, F. A., B. M. Parker, C. L. Pylant, T. J. Smyser, A. J. Piaggio, S. L. Lance, M. P. Milleson, J. D. Austin, and S. M. Wisely. 2018. Invasion ecology of wild pigs (Sus scrofa) in Florida, USA: the role of humans in the expansion and colonization of an invasive wild ungulate. Biological Invasions:1–16.

Hilton, G., and J. Packham. 2003. Variation in the masting of common beech (Fagus sylvatica L.) in northern Europe over two centuries (1800–2001). Forestry 76(3):319–328.

Hodgson, D. J., and S. Townley. 2004. Methodological insight: linking management changes to population dynamic responses: the transfer function of a projection matrix perturbation. Journal of Applied Ecology 41(6):1155–1161.

Hufbauer, R. A., M. Szűcs, E. Kasyon, C. Youngberg, M. J. Koontz, C. Richards, T. Tuff, and B. A. Melbourne. 2015. Three types of rescue can avert extinction in a changing environment. Proceedings of the National Academy of Sciences 112(33):10557–10562.

Iles, D. T., R. Salguero-Gómez, P. B. Adler, and D. N. Koons. 2016. Linking transient dynamics and life history to biological invasion success. Journal of Ecology 104(2):399–408.

Kearney, M., and W. Porter. 2009. Mechanistic niche modelling: combining physiological and spatial data to predict species’ ranges. Ecology letters 12(4):334–350.

Keiter, D. A., F. L. Cunningham, O. E. Rhodes Jr, B. J. Irwin, and J. C. Beasley. 2016. Optimization of scat detection methods for a social ungulate, the wild pig, and experimental evaluation of factors affecting detection of scat. PloS one 11(5):e0155615.

Koenig, W. D., R. L. Mumme, W. J. Carmen, and M. T. Stanback. 1994. Acorn production by oaks in central coastal California: variation within and among years. Ecology 75(1):99–109.

Kutner, M. H., C. J. Nachtsheim, J. Neter, and W. Li. 2005. Applied linear statistical models. McGraw-Hill Irwin Boston.

Lewis, J. S., M. L. Farnsworth, C. L. Burdett, D. M. Theobald, M. Gray, and R. S. Miller. 2017. Biotic and abiotic factors predicting the global distribution and population density of an invasive large mammal. Sci Rep 7:44152. doi: 10.1038/srep44152

Lustig, A., S. P. Worner, J. P. Pitt, C. Doscher, D. B. Stouffer, and S. D. Senay. 2017. A modeling framework for the establishment and spread of invasive species in heterogeneous environments. Ecology and evolution 7(20):8338–8348.

Mack, R. N., D. Simberloff, W. Mark Lonsdale, H. Evans, M. Clout, and F. A. Bazzaz. 2000. Biotic invasions: causes, epidemiology, global consequences, and control. Ecological applications 10(3):689–710.

Mayer, J., and I. Brisbin. 1991. Wild pigs in the United States: their life history, morphology and current status. University of Georgia Press. Athens, Georgia, USA.

McClure, M. L., C. L. Burdett, M. L. Farnsworth, M. W. Lutman, D. M. Theobald, P. D. Riggs, D. A. Grear, and R. S. Miller. 2015. Modeling and mapping the probability of occurrence of invasive wild pigs across the contiguous United States. PLoS One 10(8):e0133771. doi: 10.1371/journal.pone.0133771

McClure, M. L., C. L. Burdett, M. L. Farnsworth, S. J. Sweeney, and R. S. Miller. 2018. A globally-distributed alien invasive species poses risks to United States imperiled species. Sci Rep 8(1):5331. doi: 10.1038/s41598-018-23657-z

McDonald, J. L., M. Franco, S. Townley, T. H. Ezard, K. Jelbert, and D. J. Hodgson. 2017. Divergent demographic strategies of plants in variable environments. Nature ecology & evolution 1(2):0029.

Michel, N. L., M. P. Laforge, F. M. Van Beest, and R. K. Brook. 2017. Spatiotemporal trends in Canadian domestic wild boar production and habitat predict wild pig distribution. Landscape and Urban Planning 165:30–38.

Miller, R. S. 2017. Interaction among societal and biological drivers of policy at the wildlife-agricultural interface. Colorado State University

Miller, R. S. 2020. Wildlife in the United States. Environmental Issues Today: Choices and Challenges [2 volumes]:171.

Miller, R. S., S. M. Opp, and C. T. Webb. 2018. Determinants of invasive species policy: Print media and agriculture determine US invasive wild pig policy. Ecosphere 9(8)

Miller, R. S., S. J. Sweeney, C. Slootmaker, D. A. Grear, P. A. Di Salvo, D. Kiser, and S. A. Shwiff. 2017. Cross-species transmission potential between wild pigs, livestock, poultry, wildlife, and humans: implications for disease risk management in North America. Sci Rep 7(1):7821. doi: 10.1038/s41598-017-07336-z

NASS. 2019. CropScape-cropland data layer. In: N. A. S. S. United States Department of Agriculture (ed.). United States Department of Agriculture, National Agricultural Statistics Service, Washington DC.

Ostfeld, R. S., and F. Keesing. 2000. Pulsed resources and community dynamics of consumers in terrestrial ecosystems. Trends in Ecology & Evolution 15(6):232–237.

Peterson, A. T., M. Papeş, and J. Soberón. 2015. Mechanistic and correlative models of ecological niches. European Journal of Ecology 1(2):28–38.

Quintana-Ascencio, P. F., S. M. Koontz, S. A. Smith, V. L. Sclater, A. S. David, and E. S. Menges. 2018. Predicting landscape-level distribution and abundance: Integrating demography, fire, elevation and landscape habitat configuration. Journal of Ecology 106(6):2395–2408.

Reise, K., S. Olenin, and D. W. Thieltges. 2006. Are aliens threatening European aquatic coastal ecosystems? Helgoland Marine Research 60(2):77.

Rosell, C., F. Navàs, and S. Romero. 2012. Reproduction of wild boar in a cropland and coastal wetland area: implications for management. Animal Biodiversity and Conservation 35(2):209–217.

Schley, L., and T. J. Roper. 2003. Diet of wild boar Sus scrofa in Western Europe, with particular reference to consumption of agricultural crops. Mammal review 33(1):43–56.

Simberloff, D. 2009. The role of propagule pressure in biological invasions. Annual Review of Ecology, Evolution, and Systematics 40:81–102.

Snow, N. P., M. A. Jarzyna, and K. C. VerCauteren. 2017. Interpreting and predicting the spread of invasive wild pigs. Journal of Applied Ecology 54(6):2022–2032.

Stott, I., S. Townley, and D. J. Hodgson. 2011. A framework for studying transient dynamics of population projection matrix models. Ecology Letters 14(9):959–970.

Tabak, M. A., M. S. Norouzzadeh, D. W. Wolfson, S. J. Sweeney, K. C. VerCauteren, N. P. Snow, J. M. Halseth, P. A. Di Salvo, J. S. Lewis, and M. D. White. 2019. Machine learning to classify animal species in camera trap images: applications in ecology. Methods in Ecology and Evolution

Tabak, M. A., A. J. Piaggio, R. S. Miller, R. A. Sweitzer, and H. B. Ernest. 2016. Linking anthropogenic factors with the movement of an invasive species. In Submission

Tabak, M. A., C. T. Webb, and R. S. Miller. 2018. Propagule size and structure, life history, and environmental conditions affect establishment success of an invasive species. Scientific reports 8(1):10313.

Tremblay, R. L., J. Raventos, and J. D. Ackerman. 2015. When stable-stage equilibrium is unlikely: integrating transient population dynamics improves asymptotic methods. Annals of Botany 116(3):381–390.

Tuanmu, M. N., and W. Jetz. 2014. A global 1-km consensus land-cover product for biodiversity and ecosystem modelling. Global Ecology and Biogeography 23(9):1031–1045.

USDA. 2015. Feral Swine Damage Management: A National Approach. In: U. S. D. o. Agriculture (ed.). p 550. United States Department of Agriculture, Animal and Plant Health Inspection Service, Washington DC.

Vetter, S. G., T. Ruf, C. Bieber, and W. Arnold. 2015. What is a mild winter? Regional differences in within-species responses to climate change. PLoS One 10(7):e0132178.

