## Supplemental Methods for "Transient population dynamics drive the spread of invasive wild pigs in North America"

**INTRODUCTION**

Trees and shrubs producing a crop of hard mast (i.e., nuts) provide an important food resource for wildlife across the Northern Hemisphere. In North America, 186 different species of birds and mammals consume the acorn crop produced by oaks (*Quercus* spp.) (Van Dersal, 1940), including many valuable game species such as white-tailed deer (*Odocoileus virginianus*), wild turkeys (*Meleagris gallopavo*), black bears (*Ursus americanus*) and squirrels (McShea & Healy, 2002). In Europe, the wild pig (*Sus scrofa*) is another game animal whose population dynamics are strongly influenced by mast availability (Bieber & Ruf, 2005; Vetter *et al.*, 2015). Some of these game species have been introduced outside their native ranges for sport hunting where they have become destructive invasive species. For example, wild pigs are invasive in North America and cause an estimated $1.5 billion in damage in the United States (U.S.) alone (Pimental, 2007). The grey squirrel (*Sciurus carolinensis*) is native to North America but invasive in Europe where it is driving severe population declines in the native European red squirrel (*S. vulgaris*) (Stokstad, 2016). Given the strong effect of mast availability on populations of these and other species, there is real economic value to being able to depict the distribution of masting species for management or conservation planning.

However, incorporating information on mast availability into planning efforts requires high-resolution geospatial data depicting biodiversity metrics like relative density or species richness. But data depicting relative density have not yet been compiled for North America and, while there are geospatial data for Europe depicting the relative density of various forest-types (Brus *et al.*, 2012), additional processing would be needed to restrict those data to species that produce mast. Data depicting the species richness of mast trees are unavailable for the Northern Hemisphere. Species richness may be particularly important since masting can be asynchronous among tree species (Sork *et al.*, 1993; Liebhold *et al.*, 2004), suggesting more diverse areas provide a more stable annual food resource for wildlife (Koenig *et al.*, 1994).

The objective of this data analysis was to create geospatial datasets depicting both the relative density and species richness of mast-bearing species across the Northern Hemisphere. We created these datasets by compiling and reprocessing existing data depicting the occurrence and distribution of mast-producing species across Europe and North America.

**METHODS**

Europe

*Relative density*. – We used spatial data depicting the proportional land area for various European mast species to map the relative density for Europe (Tröltzsch *et al.*, 2009; Brus *et al.*, 2012) (available from <http://www.efi.int/portal/virtual_library/information_services/mapping_services/tree_species_maps_for_european_forests/>). These data were the output from statistical models consisting of a large dataset of plot-level forest inventory data as a response variable, and various environmental conditions as predictor variables (Brus *et al.*, 2012). These data, which had a 1.0 km^2^ resolution, were compiled by genus, so we created our relative-density map by summing the distributional data for all mast-producing genera in ArcGIS 10.3 (Environmental Systems Research Institute, 2014).

*Species richness*. – We measured species richness in Europe with a recently published dataset of digitized vector-based polygon maps depicting the geographic ranges of masting species (Caudullo *et al.*, 2017) (available from <https://data.mendeley.com/datasets/hr5h2hcgg4/2>). These data measured the species’ geographic ranges as extents of occurrence rather than the actual areas of occupancy (Gaston & Fuller, 2009). The use of extent-of-occurrence data is common in studies measuring species richness (Jenkins *et al.*, 2013).

We converted the vector-based polygon data for each species into raster data with a 1.0 km^2^ resolution (same resolution as the European relative-density dataset) in ArcGIS 10.3 (Environmental Systems Research Institute, 2014). A value of 1 was assigned each 1.0 km^2^ cell where the species has been documented occurring, and values of 0 were assigned to those cells lacking occurrence records. We created our species-richness map by summing this binary distributional data for all masting species.

North America

*Relative density.* – We measured relative density of masting species in the U.S. using a dataset that, similar to the data we used in Europe, was derived from statistical models using a large dataset of plot-level inventory data as a response variable, and various environmental factors as predictor variables (Wilson *et al.*, 2012) (available from https://www.fs.usda.gov/rds/archive/Product/RDS-2013-0013). This dataset, which used the Forest Inventory and Analysis (FIA) dataset compiled by the U.S. Forest Service (McRoberts *et al.*, 2005), consisted of 0.0625 km^2^ resolution raster data depicting the live basal area of 324 tree species. We identified the species bearing hard mast, summed the maps for those species, and estimated the total basal area of mast-producing species in the U.S.

To place this U.S. dataset into the same proportional-area units of the European dataset (Brus *et al.*, 2012), we transformed the basal-area units into proportions by dividing the total basal area of mast species by the total basal area of all tree species and multiplying that proportion by 100 (Tröltzsch *et al.*, 2009; Brus *et al.*, 2012). We then aggregated the 0.0625 km^2^ resolution of the U.S. data to match the 1.0 km^2^ resolution of our European data by taking the mean value from the sixteen 250 m input cells comprising an aggregated 1.0 km output cell.

*Species richness.* – We compiled new range maps for individual mast-producing species with the same dataset used to compile relative-density values for the U.S. (Wilson *et al.*, 2012) (available from https://www.fs.usda.gov/rds/archive/Product/RDS-2013-0013). We set our minimum mapping resolution by converting the raw basal-area data into binary presence-absence data using a threshold basal-area value of 0.005. We summed these binary maps to estimate the species richness of mast-producing species in the U.S. Finally, we aggregated the 0.0625 km^2^ input cell into a 1.0 km^2^ resolution raster. Unlike Europe, this dataset depicts more finely grained richness patterns as the underlying data represents the area of occupancy, rather than the coarser extent of occurrence (Gaston & Fuller, 2009). All spatial analyses for the U.S. were also done with ArcGIS 10.3 (Environmental Systems Research Institute, 2014).

**RESULTS**

Europe

*Relative density*. – The input data used to measure relative density for Europe consisted of data depicting the distribution of the European beech (*Fagus sylvatica*), sweet chestnut (*Castanea* *sativa*), and oak (*Quercus* spp.) (Brus et al. 2012). The map depicting proportional area of mast species in Europe indicated high relative densities in southern Portugal and central Germany, and in the regions with high species-richness values (northern Spain, France, Italy, and the Balkan Peninsula) (Figure 1).

*Species richness*. – The species-richness dataset contained 10 oak species (*Quercus* spp.), one chestnut species (*C.* *sativa*), and one beech species (*F. sylvatica*) (Table 1). Hotspots of masting tree species richness in Europe included northern Spain, France, Italy, and countries comprising the Balkan Peninsula, while no masting species occurred in central and northern portions of the Scandinavian countries Norway, Sweden, and Finland (Figure 1).

North America

*Relative density*. – The U.S. dataset contained 39 oak species, 13 hickory species (*Carya* spp.), five walnut species (*Juglans spp.*), and one beech species (*F. grandifolia*) (see Appendix S1). A map depicting proportional area of mast species in the U.S. indicates relatively widespread occurrence throughout the eastern U.S. with hotspots of relative density in the southern east-central U.S. (e.g., Arkansas, Missouri, Oklahoma), Appalachian Mountains, with other hotspots in the western and southwestern U.S. (e.g., Texas, California) (Figure 2). Our relative density data for North America only depict conditions for the U.S. as compiled forest inventory data for Canada and Mexico were unavailable.

*Species richness*. – The U.S. dataset also contained 39 oak species (*Quercus* spp.), 13 hickory species (*Carya* spp.), five walnut species (*Juglans spp.*), and one beech species (*Fagus grandifolia*) (Table 2). The map for these species identifies much of the eastern U.S. as a hotspot of species richness for mast-bearing species, with particularly high diversity values occurring in Alabama, Arkansas, Kentucky, Tennessee, Mississippi, and Missouri (Figure 2).

**DISCUSSION**

We found sufficient data to map both of our metrics in Europe and the U.S. In Eurasia, data were unavailable for Russia and Asia. In North America, a lack of previously compiled map data from statistical models precluded our ability to map either metric in Canada and Mexico. The lack of geographic distribution data in these regions is a common condition known as the Wallacean shortfall (Riddle *et al.*, 2011), which frequently impacts our ability to compile comprehensive biodiversity informatics at global scales.

The species richness of mast species was higher in North America than Europe. However, the higher species richness in North America does support a previous study that identified large portions of the U.S. as suitable habitat for invasive species such as wild pigs that use mast as a food source (McClure *et al.*, 2015). Since humans often introduce wild pigs into novel environments (Tabak *et al.*, 2017), a better understanding of the distribution of masting species could help predict population growth rates and probability of establishment following introduction, potentially reducing the costs of population-management programs for invasive wild pigs (Tabak *et al.*, 2018). More generally, our data could further enhance our understanding of how easily vertebrate species from Europe can become established in North America and vice-versa (Jeschke & Strayer, 2005).

**REFERENCES**

Bieber, C. & Ruf, T. (2005) Population dynamics in wild boar Sus scrofa: ecology, elasticity of growth rate and implications for the management of pulsed resource consumers. *Journal of Applied Ecology*, **42**, 1203-1213.

Brus, D.J., Hengeveld, G.M., Walvoort, D.J.J., Goedhart, P.W., Heidema, A.H., Nabuurs, G.J. & Gunia, K. (2012) Statistical mapping of tree species over Europe. *European Journal of Forest Research*, **131**, 145-157.

Caudullo, G., Welk, E. & San-Miguel-Ayanz, J. (2017) Chorological maps for the main European woody species. *Data in Brief*, **12**, 662-666.

Environmental Systems Research Institute (2014) *ArcGIS 10.3*. ESRI.

Gaston, K.J. & Fuller, R.A. (2009) The sizes of species’ geographic ranges. *Journal of Applied Ecology*, **46**, 1-9.

Jenkins, C.N., Pimm, S.L. & Joppa, L.N. (2013) Global patterns of terrestrial vertebrate diversity and conservation. *Proceedings of the National Academy of Sciences*, **110**, E2602-E2610.

Jeschke, J.M. & Strayer, D.L. (2005) Invasion success of vertebrates in Europe and North America. *Proceedings of the National Academy of Sciences of the United States of America*, **102**, 7198-7202.

Koenig, W.D., Mumme, R.L., Carmen, W.J. & Stanback, M.T. (1994) Acorn production by oaks in central coastal California: variation within and among years. *Ecology*, **75**, 99-109.

Liebhold, A., Sork, V., Peltonen, M., Koenig, W., Bjørnstad, O.N., Westfall, R., Elkinton, J. & Knops, J.M.H. (2004) Within-population spatial synchrony in mast seeding of North American oaks. *Oikos*, **104**, 156-164.

McClure, M.L., Burdett, C.L., Farnsworth, M.L., Lutman, M.W., Theobald, D.M., Riggs, P.D., Grear, D.A. & Miller, R.S. (2015) Modeling and mapping the probability of occurrence of invasive wild pigs across the contiguous United States. *PLoS ONE*, **10**, e0133771.

McRoberts, R.E., Bechtold, W.A., Patterson, P.L., Scott, C.T. & Reams, G.A. (2005) The enhanced Forest Inventory and Analysis program of the USDA Forest Service: historical perspective and announcement of statistical documentation. *Journal of Forestry*, **103**, 304-308.

McShea, W.J. & Healy, W.M. (2002) *Oak forest ecosystems: ecology and management for wildlife*. Johns Hopkins University Press, Baltimore, Maryland, USA.

Ostfeld, R.S., Canham, C.D., Oggenfuss, K., Winchcombe, R.J. & Keesing, F. (2006) Climate, deer, rodents, and acorns as determinants of variation in Lyme-disease risk. *PLOS Biology*, **4**, e145.

Pimental, D. (2007) Environmental and economic costs of vertebrate species invasions into the United States. *Manging vertebrate invasive species: proceedings of an international symposium.* (ed by G.W. Witmer, W.C. Pitt, and K.A. Fagerstone). Fort Collins, Colorado, USA.

Riddle, B.R., Ladle, R.J., Lourie, S.A. & Whittaker, R.J. (2011) Basic biogeography: estimating biodiversity and mapping nature. *Conservation Biogeography* (ed. by R.J. Ladle, And R.J. Whittaker), pp. 47-92. Blackwell Publishing, Ltd., John Wiley and Sons, Ltd., West Sussex, UK.

Rodríguez-Correa, H., Oyama, K., MacGregor-Fors, I. & González-Rodríguez, A. (2015) How are oaks distributed in the Neotropics? A perspective from species turnover, areas of endemism, and climatic niches. *International Journal of Plant Sciences*, **176**, 222-231.

Sork, V.L., Bramble, J. & Sexton, O. (1993) Ecology of mast-fruiting in three species of North American deciduous oaks. *Ecology*, **74**, 528-541.

Stokstad, E. (2016) Red squirrels rising. *Science*, **352**, 1268-1271.

Tabak, M.A., Webb, C.T. & Miller, R.S. (2018) Propagule size and structure, life history, and environmental conditions affect establishment success of an invasive species. *Scientific Reports*, **8**, 10313.

Tabak, M.A., Piaggio, A.J., Miller, R.S., Sweitzer, R.A. & Ernest, H.B. (2017) Anthropogenic factors predict movement of an invasive species. *Ecosphere*, **8**, e01844.

Tröltzsch, K., Van Brusselen, J. & Schuck, A. (2009) Spatial occurrence of major tree species groups in Europe derived from multiple data sources. *Forest Ecology and Management*, **257**, 294-302.

Van Dersal, W.R. (1940) Utilization of oaks by birds and mammals. *The Journal of Wildlife Management*, **4**, 404-428.

Vetter, S.G., Ruf, T., Bieber, C. & Arnold, W. (2015) What is a mild winter? Regional differences in within-species responses to climate change. *PLoS ONE*, **10**, e0132178.

Wilson, B.T., Lister, A.J. & Riemann, R.I. (2012) A nearest-neighbor imputation approach to mapping tree species over large areas using forest inventory plots and moderate resolution raster data. *Forest Ecology and Management*, **271**, 182-198.

**FIGURES**

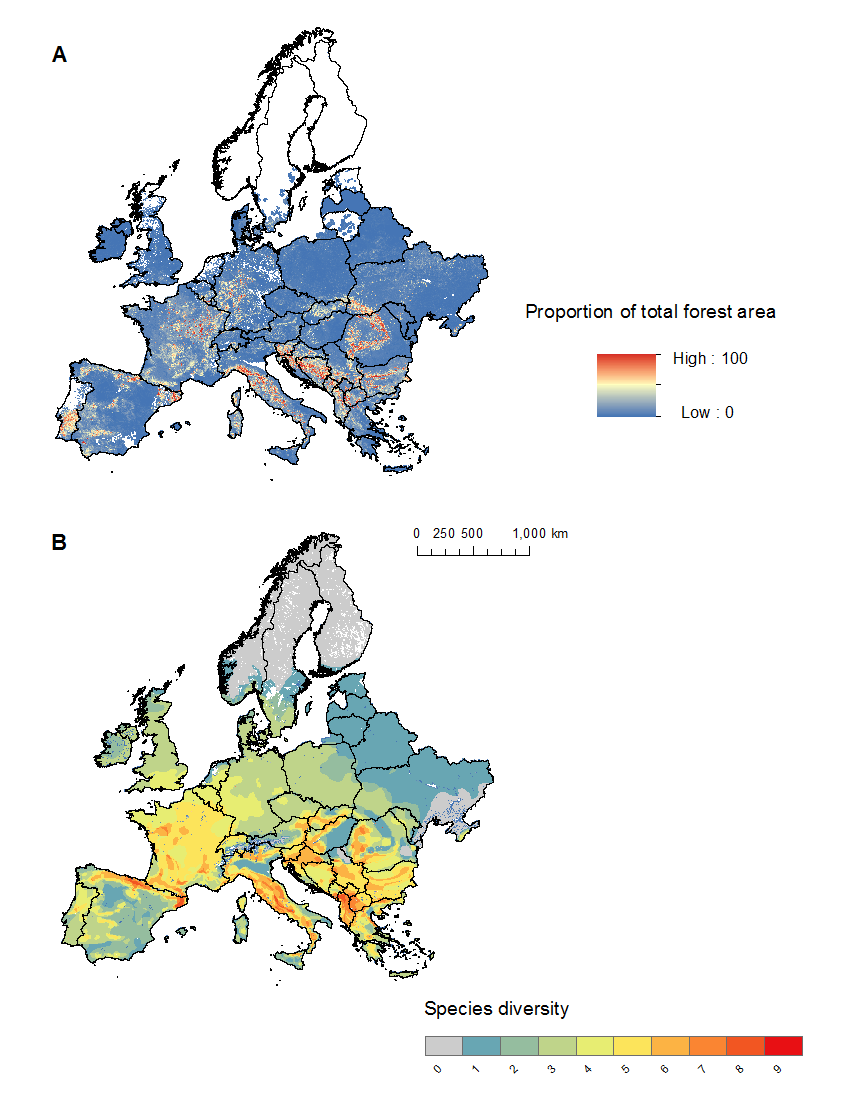

**Figure 1.** The relative density (A) and species richness (B) of species bearing hard mast in Europe. The sweet chestnut (*Castanea sativa*), which is primarily a cultivated species in Europe, is included in this map.

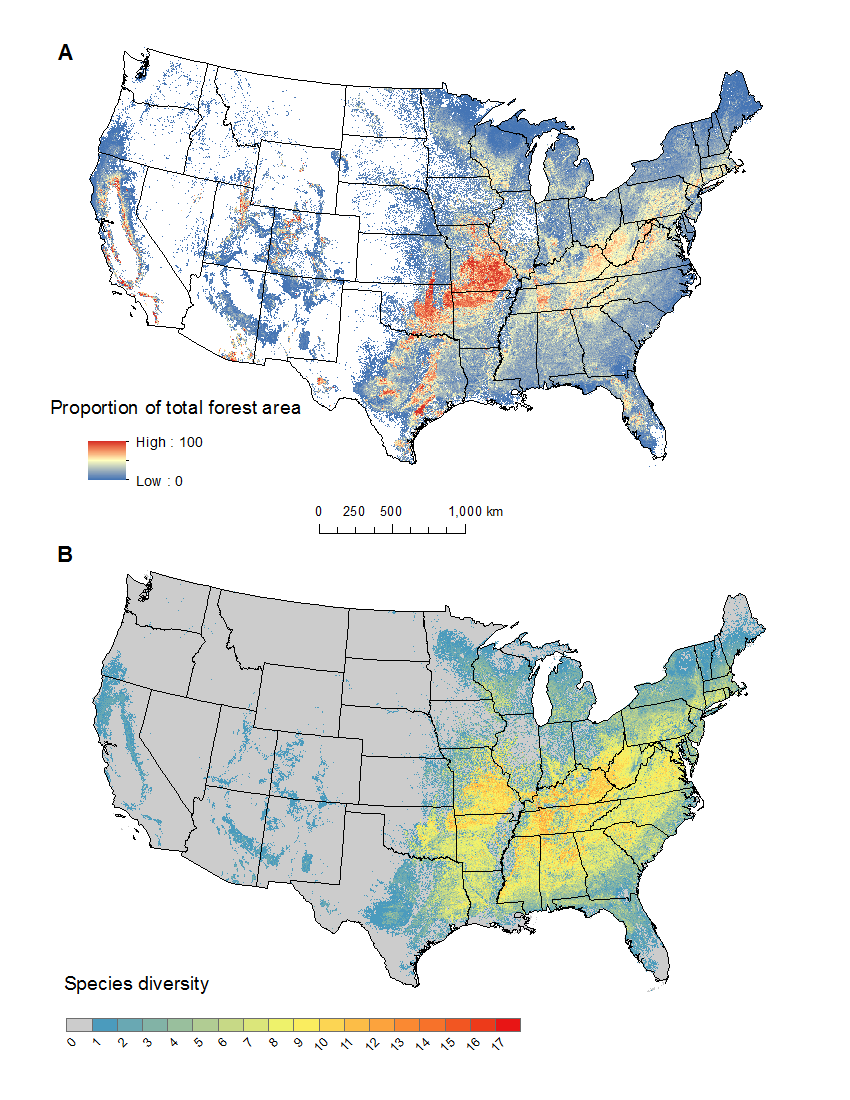

**Figure 2.** The relative density (A) and species richness (B) of species bearing hard mast in the United States (U.S.).

**TABLES**

**Table 1.** Species list for measuring species richness of mast-bearing species in Europe.

| Genus | Species | Common name |
| --- | --- | --- |
| *Castanea* | *sativa* | Sweet chestnut |
| *Fagus* | *sylvatica* | European beech |
| *Quercus* | *cerris* | Turkey oak |
| *Quercus* | *coccifera* | Kermes oak |
| *Quercus* | *frainetto* | Hungarian oak |
| *Quercus* | *ilex* | Holm oak |
| *Quercus* | *petraea* | Sessile oak |
| *Quercus* | *pubescens* | Downy oak |
| *Quercus* | *pyrenaica* | Pyrenean oak |
| *Quercus* | *robur* | Pedunculate oak |
| *Quercus* | *suber* | Cork oak |
| *Quercus* | *trojana* | Macedonian oak |

**Table 2.** Species list for measuring relative density and species of mast-bearing species in the United States.

| Genus | Species | Common name |
| --- | --- | --- |
| *Carya* | *alba* | mockernut hickory |
| *Carya* | *aquatica* | water hickory |
| *Carya* | *carolinae-septentrionalis* | southern shagbark hickory |
| *Carya* | *cordiformis* | bitternut hickory |
| *Carya* | *glabra* | pignut hickory |
| *Carya* | *illinoinensis* | pecan |
| *Carya* | *laciniosa* | shellbark hickory |
| *Carya* | *myristiciformis* | nutmeg hickory |
| *Carya* | *ovalis* | red hickory |
| *Carya* | *ovata* | shagbark hickory |
| *Carya* | *pallida* | sand hickory |
| *Carya* | spp. | hickory spp. |
| *Carya* | *texana* | black hickory |
| *Fagus* | *grandifolia* | American beech |
| *Juglans* | *cinerea* | butternut |
| *Juglans* | *major* | Arizona walnut |
| *Juglans* | *microcarpa* | Texas walnut |
| *Juglans* | *nigra* | black walnut |
| *Juglans* | spp. | walnut spp. |
| *Quercus* | *alba* | white oak |
| *Quercus* | *bicolor* | swamp white oak |
| *Quercus* | *chrysolepis* | canyon live oak |
| *Quercus* | *coccinea* | scarlet oak |
| *Quercus* | *ellipsoidalis* | northern pin oak |
| *Quercus* | *emoryi* | Emory oak |
| *Quercus* | *falcata* | southern red oak |
| *Quercus* | *gambelii* | Gambel oak |
| *Quercus* | *grisea* | gray oak |
| *Quercus* | *hypoleucoides* | silverleaf oak |
| *Quercus* | *ilicifolia* | scrub oak or bear oak |
| *Quercus* | *imbricaria* | shingle oak |
| *Quercus* | *incana* | bluejack oak |
| *Quercus* | *kelloggii* | California black oak |
| *Quercus* | *laevis* | turkey oak |
| *Quercus* | *laurifolia* | laurel oak |
| *Quercus* | *lyrata* | overcup oak |
| *Quercus* | *macrocarpa* | bur oak |
| *Quercus* | *margarettiae* | dwarf post oak |
| *Quercus* | *marilandica* | blackjack oak |
| *Quercus* | *michauxii* | swamp chestnut oak |
| *Quercus* | *minima* | dwarf live oak |
| *Quercus* | *muehlenbergii* | chinkapin oak |
| *Quercus* | *nigra* | water oak |
| *Quercus* | *oglethorpensis* | Oglethorpe oak |
| *Quercus* | *pagoda* | cherrybark oak |
| *Quercus* | *palustris* | pin oak |
| *Quercus* | *phellos* | willow oak |
| *Quercus* | *prinoides* | dwarf chinkapin oak |
| *Quercus* | *prinus* | chestnut oak |
| *Quercus* | *rubra* | northern red oak |
| *Quercus* | *shumardii* | Shumard oak |
| *Quercus* | *similis* | Delta post oak |
| *Quercus* | *sinuata* | Durand oak |
| *Quercus* | spp. | oak spp |
| *Quercus* | *stellata* | post oak |
| *Quercus* | *texana* | Texas red oak |
| *Quercus* | *velutina* | black oak |
| *Quercus* | *virginiana* | live oak |
