## Supplemental Figures and Tables for "Transient population dynamics drive the spread of invasive wild pigs in North America"

**Figure S1.** Annual watershed level spread rate (watersheds/year) for feral swine from 1982 to 2017 for three watershed scales. Red indicates positive rates of spread and blue indicates negative rates of spread (i.e. contractions in the number of occupied watersheds).

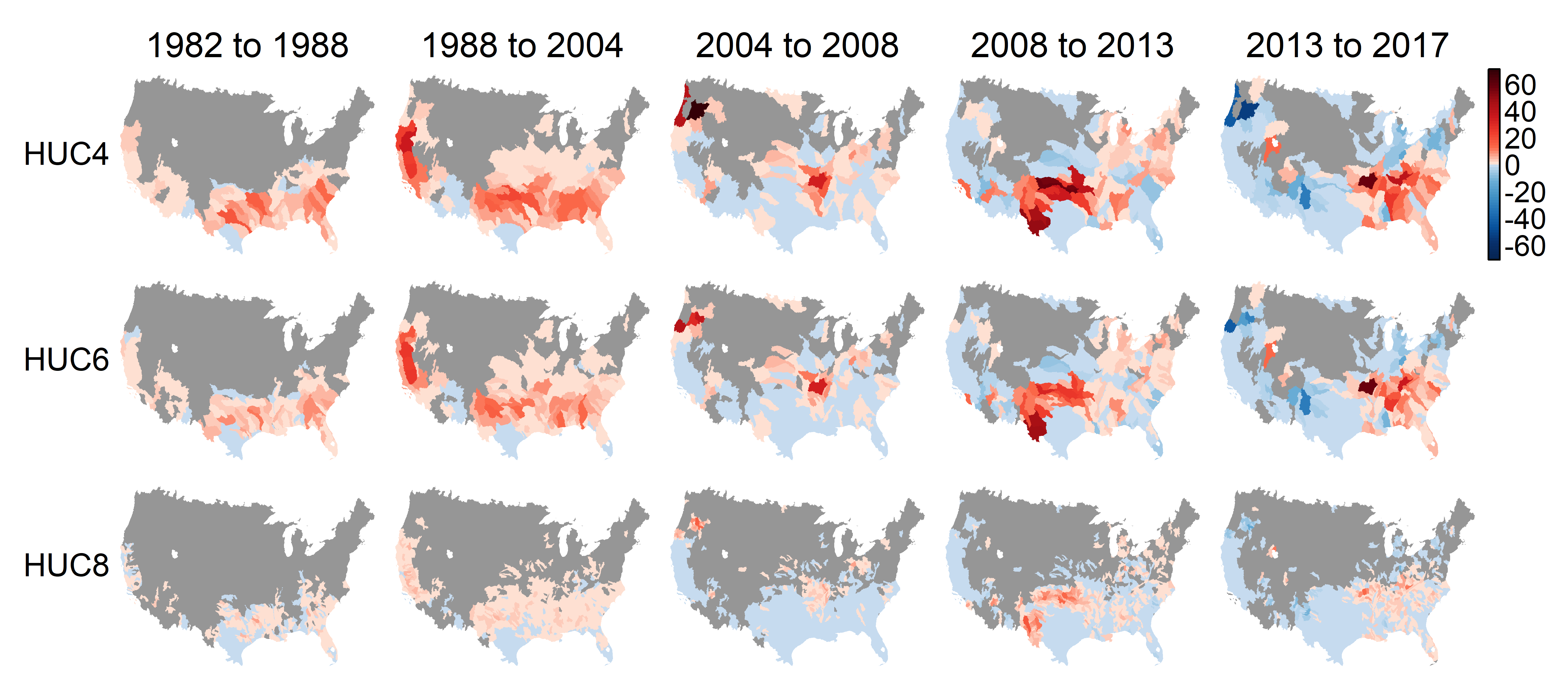

**Figure S2.** Mean annual national watershed level spread rate (watersheds/year) for feral swine from 1982 to 2017 for three watershed scales. Points represent the national level mean for the time period with the 95% confidence interval. Spread rate was significantly reduced after the APHIS national program was established.

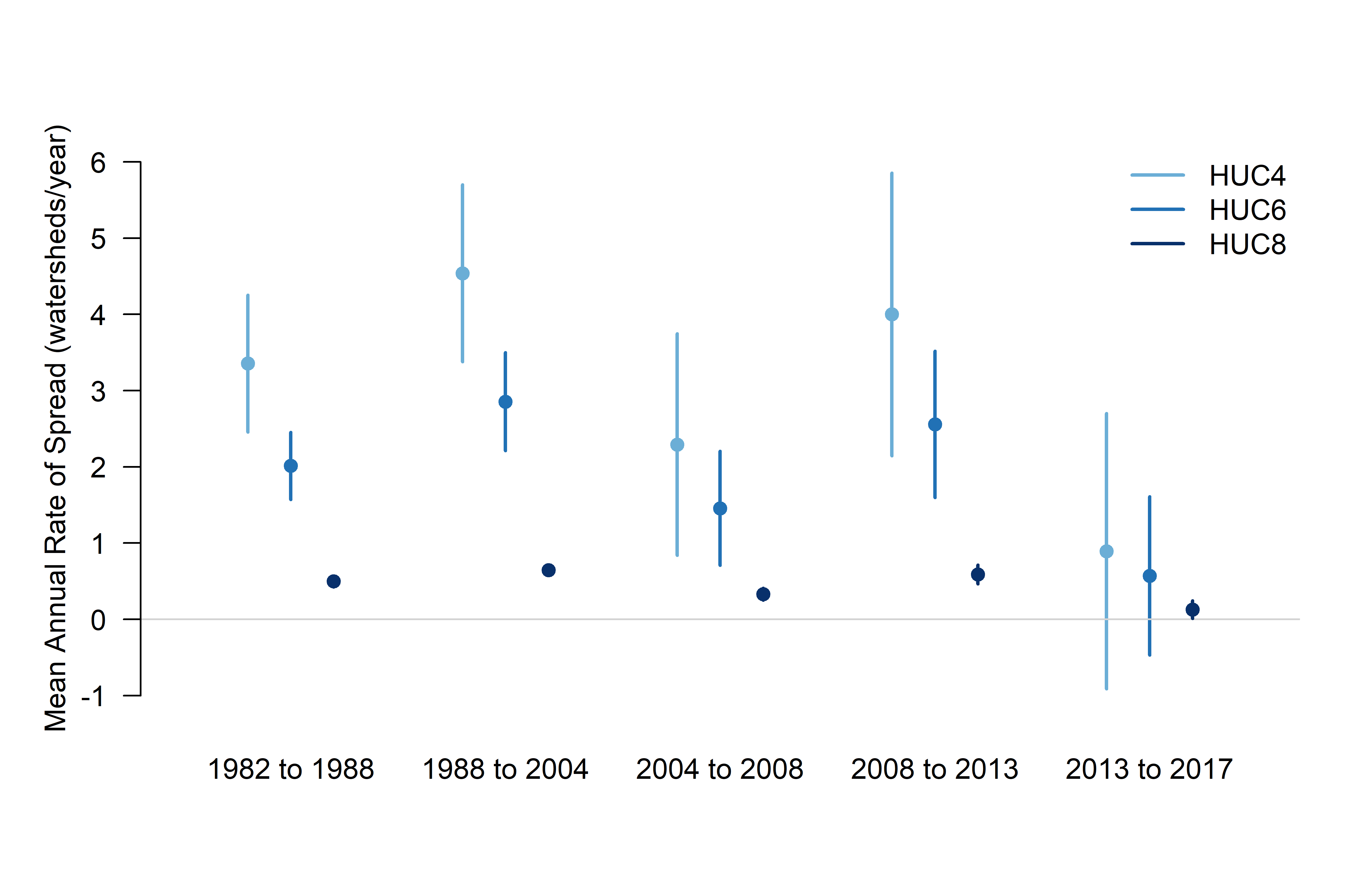

**Figure S3.** Predictor layers used in calculating population growth rates and probability of establishment.

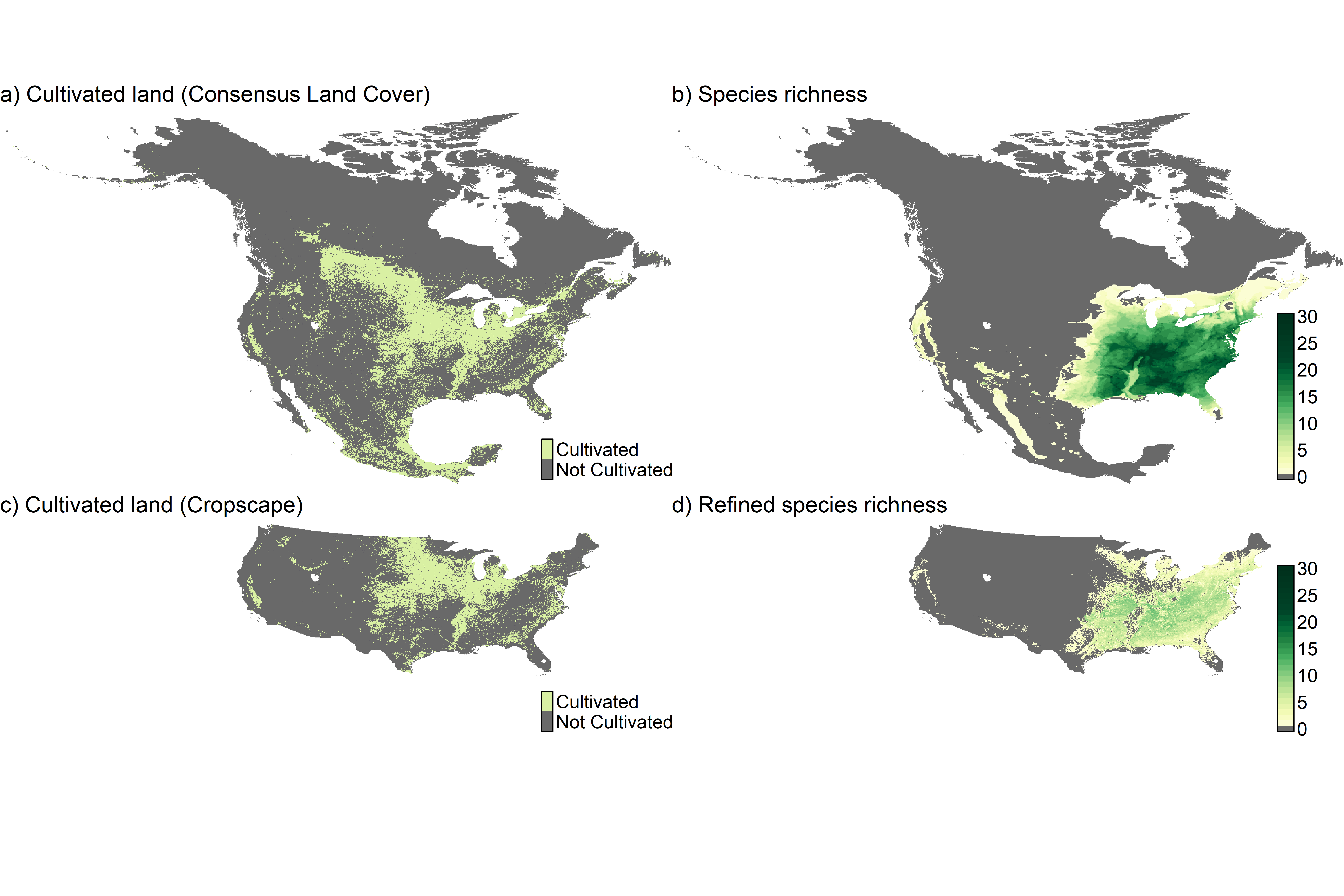

**Table S1.** Mean spread rates (watershed / year) and 95% confidence intervals for the three watershed scales used in the analysis. P-value indicates if the mean rate of spread is significantly different from the previous time period. The number of watersheds with spread rates ($\theta$) greater and less than 0 are also indicated for each period.

| **Watershed** | **Period** | **n** | **Mean** | **95% CI** | **P-Value** | $\boldsymbol{\theta}$**<0** | $\boldsymbol{\theta}$**>0** |
| --- | --- | --- | --- | --- | --- | --- | --- |
| HUC4 | 1982 to 1988 | 69 | 3.36 | (2.46, 4.25) | - | 0 | 61 |
|  | 1988 to 2004 | 112 | 4.54 | (3.38, 5.70) | 1.11 x10^-01^ | 0 | 102 |
|  | 2004 to 2008 | 122 | 2.29 | (0.84, 3.75) | 1.76 x10^-02^ | 0 | 56 |
|  | 2008 to 2013 | 142 | 4.00 | (2.15, 5.85) | 1.53 x10^-01^ | 45 | 72 |
|  | 2013 to 2017 | 135 | 0.89 | (-0.91, 2.70) | 1.83 x10^-02^ | 51 | 53 |
| HUC6 | 1982 to 1988 | 115 | 2.01 | (1.57, 2.45) | - | 0 | 91 |
|  | 1988 to 2004 | 178 | 2.86 | (2.21, 3.50) | 3.36 x10^-02^ | 0 | 155 |
|  | 2004 to 2008 | 192 | 1.46 | (0.71, 2.20) | 5.42 x10^-03^ | 0 | 70 |
|  | 2008 to 2013 | 222 | 2.56 | (1.60, 3.52) | 7.66 x10^-02^ | 61 | 113 |
|  | 2013 to 2017 | 212 | 0.57 | (-0.47, 1.61) | 5.91 x10^-03^ | 67 | 79 |
| HUC8 | 1982 to 1988 | 464 | 0.50 | (0.43, 0.57) | - | 0 | 275 |
|  | 1988 to 2004 | 788 | 0.65 | (0.59, 0.70) | 1.92 x10^-03^ | 0 | 625 |
|  | 2004 to 2008 | 846 | 0.33 | (0.25, 0.41) | 3.72 x10^-10^ | 0 | 163 |
|  | 2008 to 2013 | 963 | 0.59 | (0.47, 0.71) | 5.87 x10^-04^ | 192 | 363 |
|  | 2013 to 2017 | 948 | 0.13 | (0.01, 0.24) | 1.00 x10^-07^ | 174 | 264 |

**Table S2.** Regression coefficients for effect of lambda on watershed level spread rate.

| **Spatial Resolution** | **Period** | **Estimate** | **Std. Error** | **P-Value** |  | ***R^2^*** | **Adj. *R^2^*** |
| --- | --- | --- | --- | --- | --- | --- | --- |
| **HUC4** | 1982 to 1988 | 3.7671 | 0.1889 | 4.61 x10^-22^ | *** | 0.911 | 0.908 |
|  | 1988 to 2004 | 5.2028 | 0.3394 | 4.22 x10^-18^ | *** | 0.858 | 0.854 |
|  | 2004 to 2008 | 5.8941 | 0.5655 | 7.82 x10^-13^ | *** | 0.736 | 0.729 |
|  | 2008 to 2013 | 6.1199 | 0.4822 | 1.99 x10^-15^ | *** | 0.805 | 0.800 |
|  | 2013 to 2017 | 1.1781 | 0.7412 | 1.20 x10^-01^ |  | 0.061 | 0.037 |
|  | 1982 to 2013 | 3.7221 | 0.1849 | 3.31 x10-^22^ | *** | 0.912 | 0.910 |
|  | All Periods | 3.0594 | 0.1262 | 3.93 x10^-25^ | *** | 0.938 | 0.936 |
| **HUC6** | 1982 to 1988 | 2.5441 | 0.0964 | 1.75 x10^-26^ | *** | 0.947 | 0.946 |
|  | 1988 to 2004 | 3.4558 | 0.2155 | 9.15 x10^-19^ | *** | 0.868 | 0.865 |
|  | 2004 to 2008 | 4.3567 | 0.1942 | 6.75 x10^-24^ | *** | 0.928 | 0.926 |
|  | 2008 to 2013 | 4.3894 | 0.3321 | 5.49 x10^-16^ | *** | 0.818 | 0.813 |
|  | 2013 to 2017 | 0.6346 | 0.5132 | 2.24 x10^-01^ |  | 0.038 | 0.013 |
|  | 1982 to 2013 | 2.6398 | 0.1095 | 4.82 x10^-25^ | *** | 0.937 | 0.936 |
|  | All Periods | 2.1318 | 0.063 | 1.57 x10^-30^ | *** | 0.967 | 0.966 |
| **HUC8** | 1982 to 1988 | 0.8426 | 0.0297 | 1.18 x10^-27^ | *** | 0.954 | 0.953 |
|  | 1988 to 2004 | 0.8251 | 0.0376 | 1.48 x10^-23^ | *** | 0.925 | 0.923 |
|  | 2004 to 2008 | 1.7389 | 0.1052 | 3.31 x10^-19^ | *** | 0.875 | 0.872 |
|  | 2008 to 2013 | 1.4768 | 0.1213 | 7.44 x10^-15^ | *** | 0.792 | 0.786 |
|  | 2013 to 2017 | 0.5402 | 0.1471 | 7.19 x10^-04^ | *** | 0.257 | 0.238 |
|  | 1982 to 2013 | 0.7632 | 0.0318 | 5.62 x10^-25^ | *** | 0.937 | 0.935 |
|  | All Periods | 0.6738 | 0.0178 | 2.49 x10^-32^ | *** | 0.973 | 0.973 |
